# Label-Free Visualization and Segmentation of Endothelial Cell Mitochondria Using Holotomographic Microscopy and U-Net

**DOI:** 10.1101/2024.11.26.625487

**Authors:** Raul Michael, Tallah Modirzadeh, Tahir Bachar Issa, Patrick Jurney

## Abstract

Understanding the physiological processes underlying age-related cardiovascular disease (CVD) requires examination of endothelial cell (EC) mitochondrial networks, because mitochondrial function and adenosine triphosphate production are crucial in EC metabolism, and consequently influence CVD progression. Although current biochemical assays and immunofluorescence microscopy can reveal how mitochondrial function influences cellular metabolism, they cannot achieve live observation and tracking changes in mitochondrial networks through fusion and fission events. Holotomographic microscopy (HTM) has emerged as a promising technique for real-time, label-free visualization of ECs and their organelles, such as mitochondria. This non-destructive, non-interfering live cell imaging method offers unprecedented opportunities to observe mitochondrial network dynamics. However, because existing image processing tools based on immunofluorescence microscopy techniques are incompatible with HTM images, a machine-learning model is required. Here, we developed a model using a U-net learner with a Resnet18 encoder to identify four classes within HTM images: mitochondrial networks, cell borders, ECs, and background. This method accurately identifies mitochondrial structures and positions. With high accuracy and similarity metrics, the output image successfully provides visualization of mitochondrial networks within HTM images of ECs. This approach enables the study of mitochondrial networks and their effects, and holds promise in advancing understanding of CVD mechanisms.

## Introduction

Quantifying endothelial cell (EC) metabolism is essential for understanding the physiological and pathological processes underlying not only metabolic diseases, but also cardiovascular diseases (CVD) [1] and cancer [2]. This quantification is also critical for assessing mitochondrial function. Biochemical techniques— such as the detection of reactive oxygen species, and measurement of mitochondrial membrane potential, oxygen consumption rates, and adenosine triphosphate production—are crucial for determining the functional parameters indicating mitochondrial metabolic and bioenergetic states. Alternatively, imaging techniques may be used, such as fluorescence labeling of proteins associated with mitochondria or electron microscopy-based imaging, which requires complex preparations and is unsuitable for live cells [3]. These methods have been indispensable in elucidating the intricate mechanisms governing endothelial function and dysfunction, and may provide insight into how metabolic alterations in cells influence CVD progression, as well as the effectiveness of potential therapeutic strategies. However, traditional fluorescence-based microscopy techniques, such as wide-field fluorescence microscopy and high-resolution confocal laser scanning microscopy, have limitations including an inability to observe real-time dynamic changes in mitochondrial morphology and function, and a need for specific dye labeling, which precludes label-free analysis and introduces artifacts that might not accurately represent organelles’ unperturbed states [3].

Recently, holotomographic microscopy (HTM) has emerged as a technique enabling detailed, real-time, and label-free visualization and analysis of ECs and their intracellular components, including mitochondria [4]. HTM is a laser-based technique that measures the three-dimensional refractive index (RI) tomogram of microscopic samples, such as ECs, by using holographic imaging and inverse scattering principles. Multiple 2D holographic images of a sample are captured from various illumination angles, and a 3D RI tomogram is reconstructed through inverse solving of light scattering data [5]. HTM serves as a label-free quantitative imaging technique for microscopic phase objects, including ECs and mitochondria, with advantages including quantitative imaging capability and precise, rapid measurement over long time periods. This method facilitates the visualization and analysis of cellular membranes, subcellular organelles, and intracellular structures, without requiring exogenous labeling agents, thus providing a valuable tool for studies of EC metabolism and mitochondrial function.

A suite of tools is available for analyzing images produced using traditional fluorescence-based microscopy methods, including several tools for analyzing mitochondrial networks and their dynamics [6], [7]. However, these tools generally do not work with HTM datasets, for various technical reasons associated with the differences in RI distributions in extremely flat cells such as ECs. The cytoplasm of these cells yields RI values similar to those of the substrate, thus hindering segmentation ability [8]. The limitations of traditional methods highlight the need for innovative imaging techniques offering higher resolution and specificity, and enabling monitoring of cellular processes in real time. Novel imaging sources, combined with machine learning (ML) algorithms for image segmentation and analysis, present a promising avenue for overcoming these challenges.

ML is a scientific discipline that merges computer science and statistical methods, and enables machines to learn from experience without explicit programming. Deep learning (DL), a subset of ML, uses multiple layers of neural networks (NNs) to glean insights from data. Each layer performs mathematical operations on the input data received from the preceding layer before passing the data to the next layer. NNs are the fundamental building blocks of DL, and can adapt to any learning process with any given accuracy, after being provided sufficient data [9]. With recent advancements in computing, DL has found extensive applications in fields including image segmentation [10], [11], biology [12], computer vision [13], [14], [15], and healthcare [16], [17].

With recent advancements in computing power, ML and DL techniques have been used to achieve state-of-the-art results in image segmentation. One study [18] has used a U-Net [11] based architecture to perform segmentation of EC borders. On a test dataset of 30 images of corneal ECs of different sizes, this method achieved an AUROC of 92%, with an average F1 score of 0.86. Another study [19] has used an improved version of the U-Net model to achieve state-of-the-art results in detecting the cell boundaries of corneal ECs. Current achievements and challenges in using DL methods for image segmentation have been discussed in detail in recent articles [20], [21], [22].

Although DL has been used in image segmentation and cellular research, it has rarely been applied beyond confocal microscopy studies. HTM provides a three-dimensional framework to study cellular structures, and can achieve enhanced capabilities when it is paired with techniques such as NN. Convolutional NNs can automatically adapt to learn classifications and their associations with specific pixels. This ability to train the network to recognize pixels as certain established classes has also been explored in studies of EC mitochondrial networks, structures, and dynamics [19]. Using HTM allows the network to learn which class is associated with a desired pixel in a high-resolution, label-free image. Using HTM from video acquisition provides multiple frames without staining, thus enabling optimal training and segmentation. Such advancements have not only improved understanding of EC metabolism and its roles in diseases, but also opened new possibilities for the development of targeted therapies and interventions for CVD and related conditions.

In this work, we sought to use transfer learning techniques and U-Net based models to achieve quantification of EC mitochondrial networks through segmentation of HTM images, as follows:

1. We collected and labeled 150 HTM images for model training and testing.
2. Through transfer learning, triangular learning rates, and data augmentation techniques, we trained a U-Net model that successfully quantified ECs and sub-cellular organelles.
3. The model was tested on a new data set previously unseen by the model, and achieved an average accuracy of 95% and an average Structural Similarity Index for Measuring Image Quality (SSIM) of 0.92 in identifying the cells, cell borders, mitochondria, and background.
4. We have made the model available to the public via a web application: https://huggingface.co/spaces/itahir/SegmHTI

## Methods

### Cell Culture

Human umbilical vein endothelial cells (HUVECs) were cultured in T-75 flasks (ThermoFisher Scientific, United States) coated with collagen I (Gibco, United States). Cells were maintained in Fluorobrite Dulbecco’s Modified Eagle’s Medium (DMEM; Gibco, United States) supplemented with 5% fetal bovine serum (FBS), growth factor supplements (ATCC Primary Cell Solutions, United States), and antibiotics. The growth factors included 5 ng/mL recombinant human vascular endothelial growth factor (rhVEGF), 5 ng/mL rh epidermal growth factor (rhEGF), 5 ng/mL rh fibroblast growth factor basic (rhFGF basic), 15 ng/mL rh insulin-like growth factor (rhIGF), 10 mM L-glutamine, 0.75 units/mL heparin sulfate, and 1 μg/mL ascorbic acid. Antibiotics consisted of 1% penicillin-streptomycin and fungizone (Cytiva, United States). Cells were incubated at 37 °C in a humidified atmosphere with 5% CO₂.

When the cells grew to approximately 80% confluence, they were washed with phosphate-buffered saline (PBS) and detached using TrypLE Express (Gibco, United States). The enzymes were neutralized with 5% FBS in DMEM, the cells collected into 15 mL conical tubes, and the suspension centrifuged at 100 × g for 7 minutes to pellet the cells. The supernatant was discarded, and the cell pellet was resuspended in fresh DMEM. Cell viability and counts were determined using a trypan blue exclusion assay (Invitrogen, United States) and an automated cell counter. The cells were then seeded into collagen-coated 35 mm dishes with polymer coverslip bottoms (ibidi GmbH, Germany) for imaging.

### Holotomographic Microscopy Imaging

HTM imaging was performed using a CX-F HTM system (Nanolive SA, Switzerland) using STEVE software (Nanolive SA, Switzerland). The ibidi dishes containing the cells were placed on the microscope stage, and the x, y, and z positions were adjusted to locate the desired cells. After focusing, three-dimensional images were acquired as stacks of .vol files. These volumetric data were converted to .tiff stacks for further analysis using ImageJ software.

To enhance mitochondrial visualization, a maximum intensity projection (MIP) was applied to the raw images using a custom macro in ImageJ. The macro automated the MIP process, generating projections for each image in the stack. The resulting images were saved for subsequent analysis.

An additional set of data was collected using four-dimensional (4D) video acquisition to capture dynamic cellular behavior. Prior to recording, the stage-top incubator (Nanolive SA, Switzerland) was powered on at least 2 hours in advance to stabilize the temperature at 37 °C. One hour into the warming process, the gas mixer was activated to regulate CO₂ levels. Cells were placed on the microscope stage under the incubator to maintain optimal environmental conditions. After focusing, 4D video acquisition was initiated using the STEVE software. Prolonged video recording enabled observation of extensive cell movement. The videos produced .vol and .tiff files, which were transferred to a separate computer for post-processing. Still images were extracted from the videos for data labeling and model training.

### Data labeling and Annotation

To prepare the dataset for model training, both HTM image stacks and frames extracted from video files were used. Frames that exhibited prominent mitochondrial networks were selected to enhance the diversity of the dataset. Each selected image was manually segmented into four classes: cells, cell borders, mitochondria, and background. Images were annotated using the built-in annotation features of an Apple tablet to trace the mitochondrial networks and cell borders. The images were then imported into Adobe Photoshop for refinement, where the mitochondrial networks were precisely traced using the pencil tool. A preliminary annotated image is shown in Figure 1a, with mitochondria in green and cell borders in red. Finally, the outer part of the cell was filled with a different color from the inner part, assigning each class a designated color. An example of the final annotated image is shown in Figure 1b, where the four classes are labeled with specific colors.

**Figure 1.**
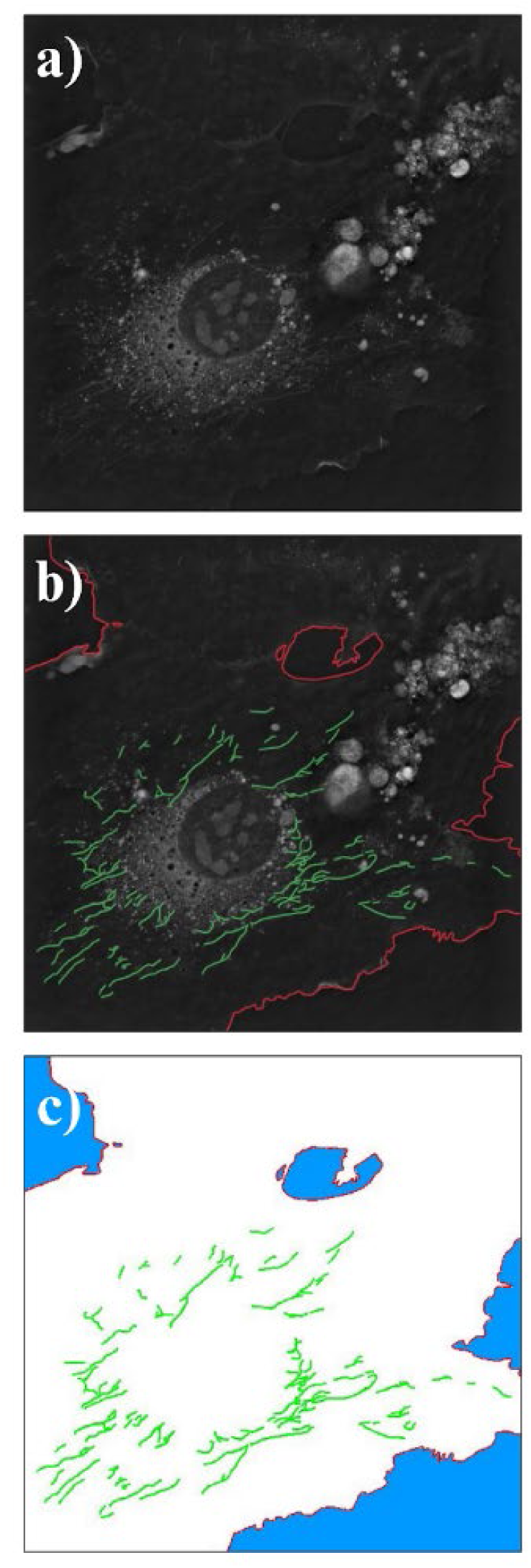
Example of the manual labeling process of holotomographic microscopy (HTM) images: (**a**) unannotated HTM image; (**b**) HTM image with labeled mitochondrial networks (green) and cell border (red); and (**c**) final annotated HTM image with the cell (white) and background (blue) labeled.

### Dataset Preparation

The resulting dataset consisted of 145 annotated images. Of these, 80% were randomly selected for model training, and the remaining 20% were used for model validation. An additional test set consisted of five unique images not included in the training or validation sets. The selected images encompassed a variety of fields of view, including entire cell circumferences, detailed cell interiors, frames rich in mitochondrial networks, and images with intricate cell border lines.

### Model Architecture

We used an improved version of the standard U-Net architecture, which was originally designed to achieve high performance in segmenting biological images in small datasets. The architecture consists of a contracting path called an encoder and a decontracting path called a decoder. The encoder uses successive sequences of convolution layers, followed by rectified linear unit (ReLU), batch normalization, and pooling of layers to map the input image to a lower-dimensional space while extracting important features. The decoder reconstructs a transformed version of the input image by combining concatenation and deconvolution techniques with the features extracted by the encoder.

To enhance model convergence and significantly improve performance, we used a U-Net architecture with a pretrained ResNet18 [23] encoder, referred to as Dynamic U-Net [24], [25]. This pretrained encoder has been trained on more than 1 million images in the ImageNet dataset. Using a pretrained encoder has been shown to accelerate model convergence and significantly enhance model performance. For assessing model performance during model training for our multiclass classification problem, we used a weighted cross entropy loss function given by:

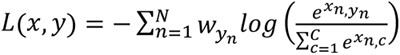

where *N* is the batch size, 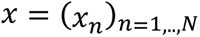 is an *N* × *C* matrix with 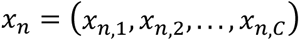 as the model prediction for batch 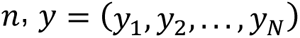 is the target vector of the batch, *C* is the number of classes, and 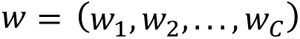 is the weight vector.

Finally, for the model parameter updates, we used the newly introduced, state-of-the art DL optimizer Ranger [26] with the Mish activation function, as previously suggested [27], [28].

The detailed architecture of the model used is shown in Figure 2.

**Figure 2.**
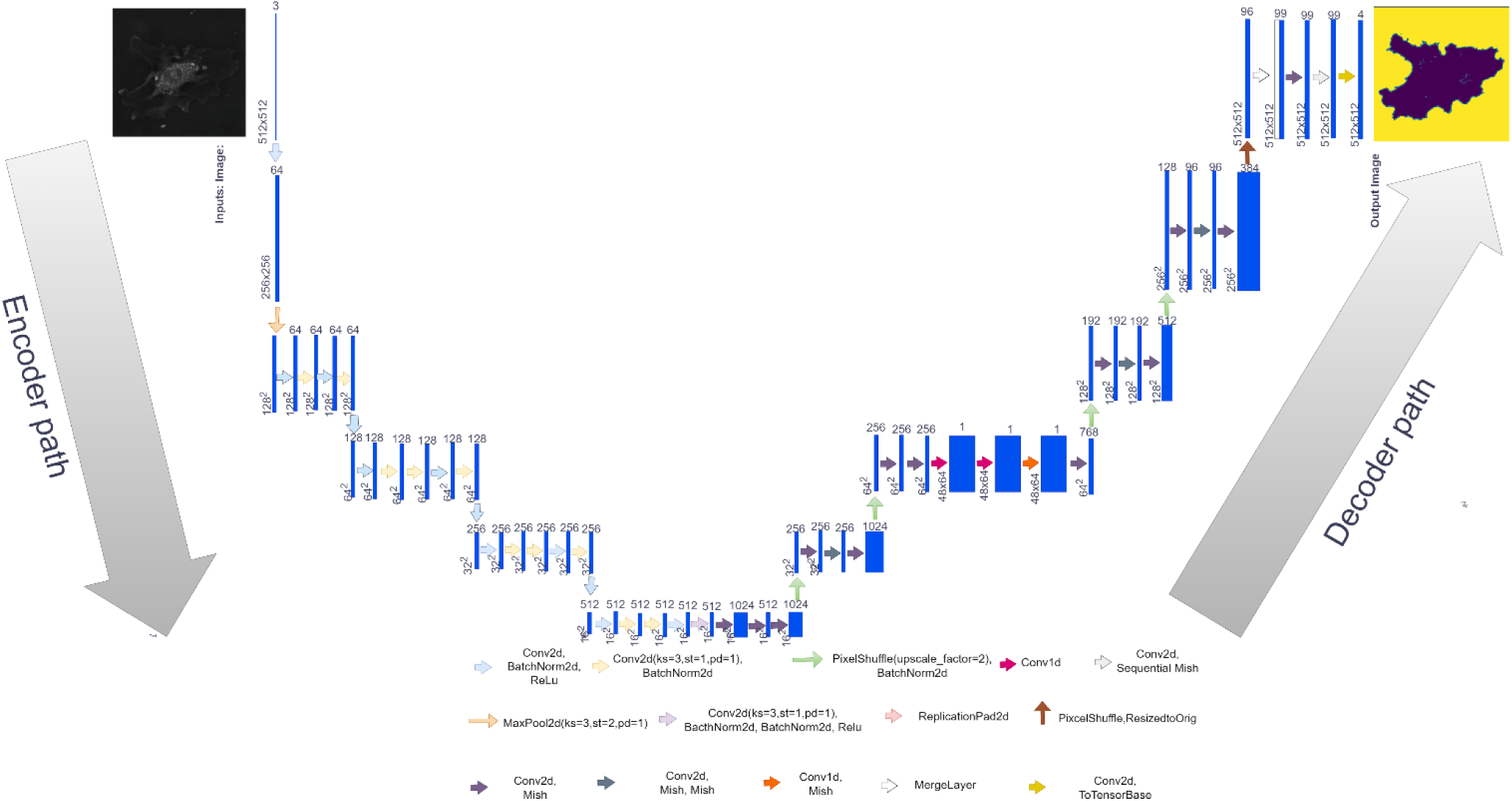
Dynamic U-Net architecture used to create the machine learning algorithm, beginning with the raw image and outputting the predicted image.

### Model Training

The U-Net learner was trained through an iterative approach in FastAI (Figure 3). For increasing data size and diversity, images were randomly augmented with a rotation of 20° or a zoom factor of 1.2. After augmentation, the model was trained for 100 epochs with an initial learning rate of 0.0003, a weight decay of 0.01, and a batch size of 5. A built-in FastAI function automatically reduced the learning rate when plateauing was observed in the training and validation results. A custom accuracy function was written, which calculated segmentation accuracy according to the model’s identification of every class excluding the background. This accuracy function, paired with validation loss, was used to identify the highest-quality trained model. Additionally, qualitative analysis of the validation results was used to determine whether overfitting was present. If overfitting was observed, the weighted loss function was adjusted to address the overfitting of specific classes. After the model with the highest accuracy was identified, the test images were inputted to determine the model’s accuracy in segmenting real-world data.

**Figure 3.**
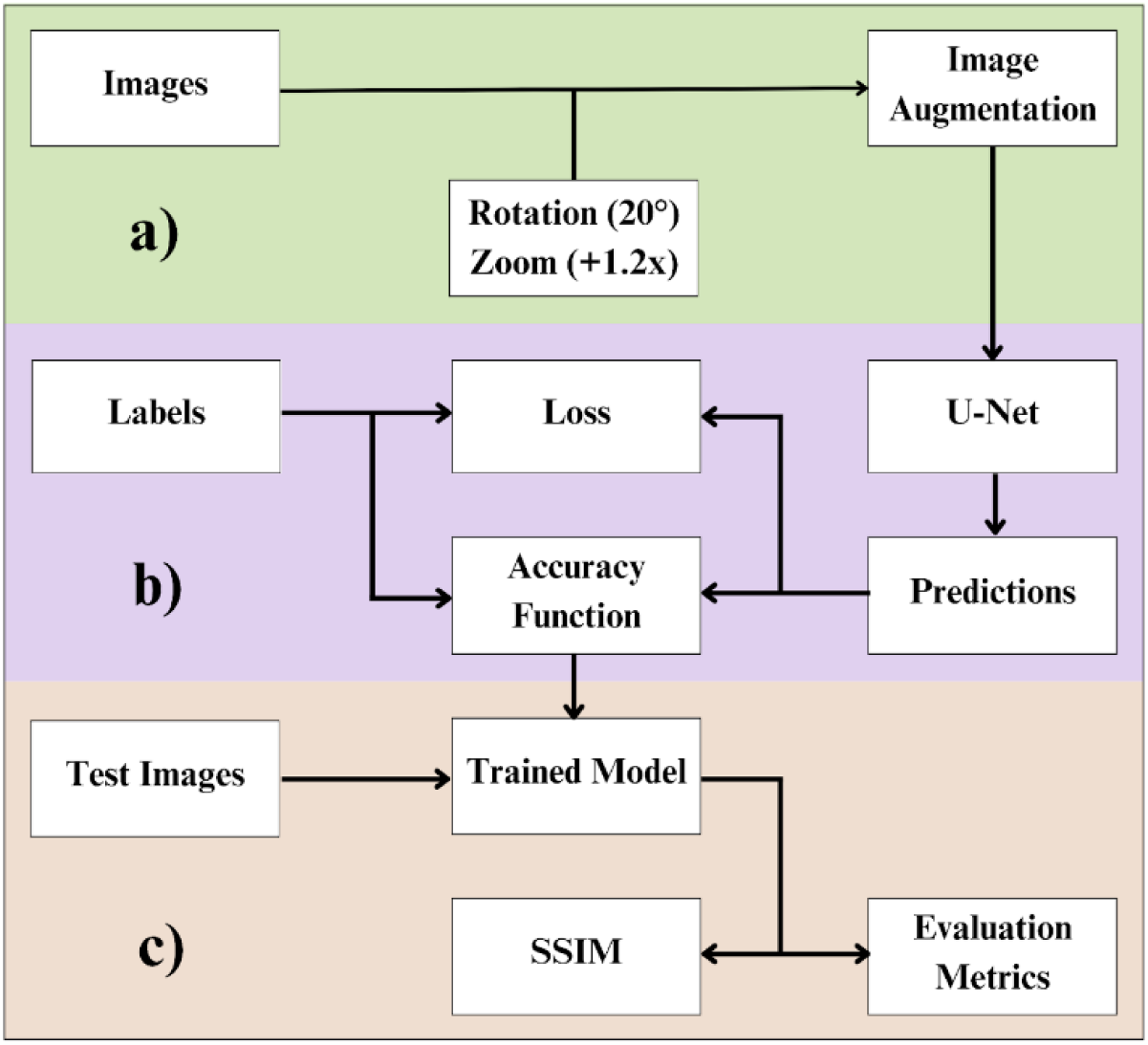
Machine learning training schematic showing the (**a**) input data process and (**b**) iterative model training process, wherein training loss, validation loss, and an accuracy function are used as training metrics, and (**c**) testing and metrics used for model evaluation.

**Figure 4.**
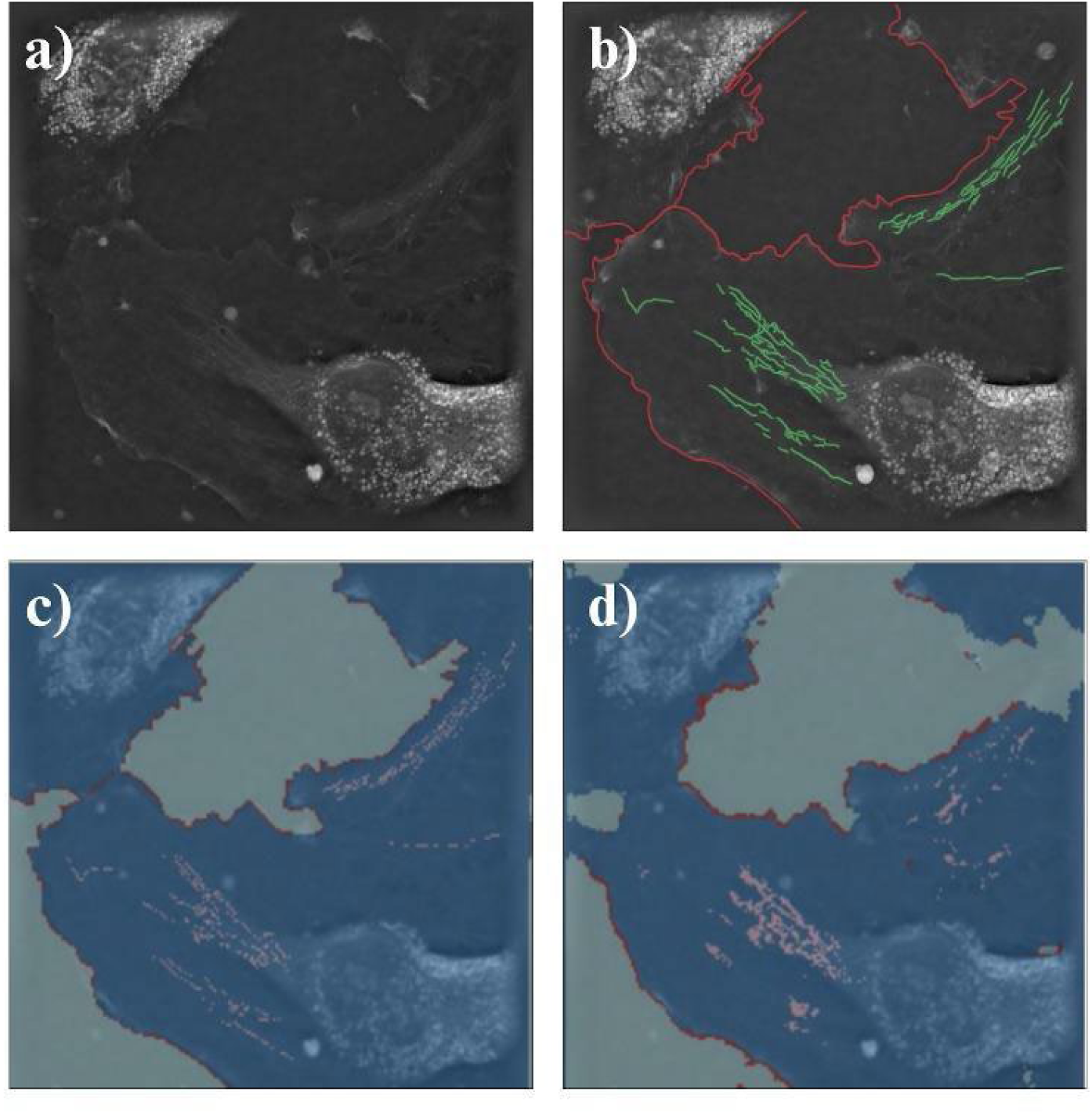
Qualitative representation of the image segmentation pipeline: (**a**) raw, unannotated image, (**b**) ground truth, (**c**) target mask, and (**d**) model output.

### Model Evaluation

To determine how accurately the model segmented the validation and the test sets, which had imbalanced classes, we calculated precision, recall, and F1 score metrics. Additionally, we determined the SSIM to compare the model output with the corresponding ground truth images.

Precision, a commonly used metric, indicates the ratio between the model-predicted number of positive samples and the true number of positive samples.

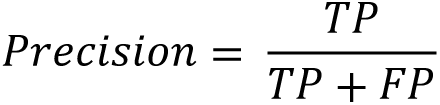

This metric is used to analyze the accuracy of positive predictions by accounting for true positives (TP), which are the correctly predicted positive instances, and false positives (FP), which are negative instances incorrectly predicted as positive. This metric is important in discerning the model’s ability to make false positive predictions, and it provides insight into the quality of the model’s positive predictions.

Similarly, recall is a commonly used metric indicating the proportion of actual positive instances correctly predicted by the model.

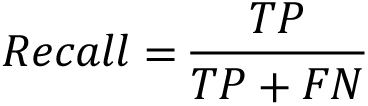

The recall metric measures the model’s ability to identify all relevant positive examples. It finds the proportion of TP to the sum of TP and false negatives (FN), which are positive instances incorrectly predicted as negative.

This metric determines the model’s ability to make false negative predictions and highlights the quantity of the model’s positive predictions and its accuracy in those predictions.

The F1 score, the harmonic mean of precision and recall, enables overall evaluation of the model’s ability to predict positive instances, while accounting for both FP and FN.

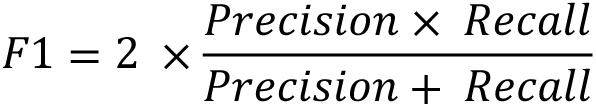

The F1 score indicates the balance between the precision and recall metrics [20], and is especially helpful when class imbalances are present, because accuracy functions can be misleading when majority and minority classes are present.

SSIM is a metric used to compare the output images with the ground truth images, to quantify the similarity between the identification of classes in the two images.

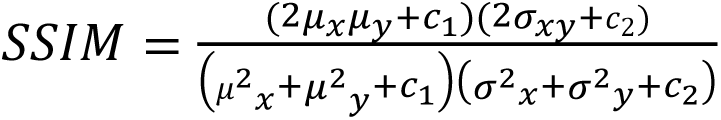

In this analysis, *x* and *y* denote the reference image and the input image, respectively. *u_x_* and *u_y_* represent the mean pixel intensities of *x* and *y*, quantifying the luminance in each image. The terms *σ_x_* and *σ_x_* correspond to the variances of the pixel intensities, thus providing a measure of contrast for the reference and input images. The covariance *σ_xy_* captures the degree of structural correlation between the images, reflecting the extent to which their spatial structures vary in alignment.

The SSIM metric evaluates luminance, contrast, and structure to assess image similarity [29]. Each factor—luminance (brightness of each pixel), contrast (pixel value variation), and structure (pixel correlation across images)—is assessed independently and is then combined into a single metric. This SSIM score enables a comprehensive comparison of the input image to the ground truth, thus serving as an accuracy metric incorporating the key attributes of each image.

Both the validation set and the test set achieved precision, recall, and F1 scores above 0.9, thereby indicating highly accurate segmentation Thus, the model identified each class with minimal error, and minimal instances of FP and FN. Each test image showed high accuracy and structural similarity measurements (Figure 5), with an average SSIM score above 0.9. Although more interconnected network fragments cannot be visually discerned, the output images show substantial promise in outlining the overall shape of the cell and mitochondrial networks.

**Figure 5.**
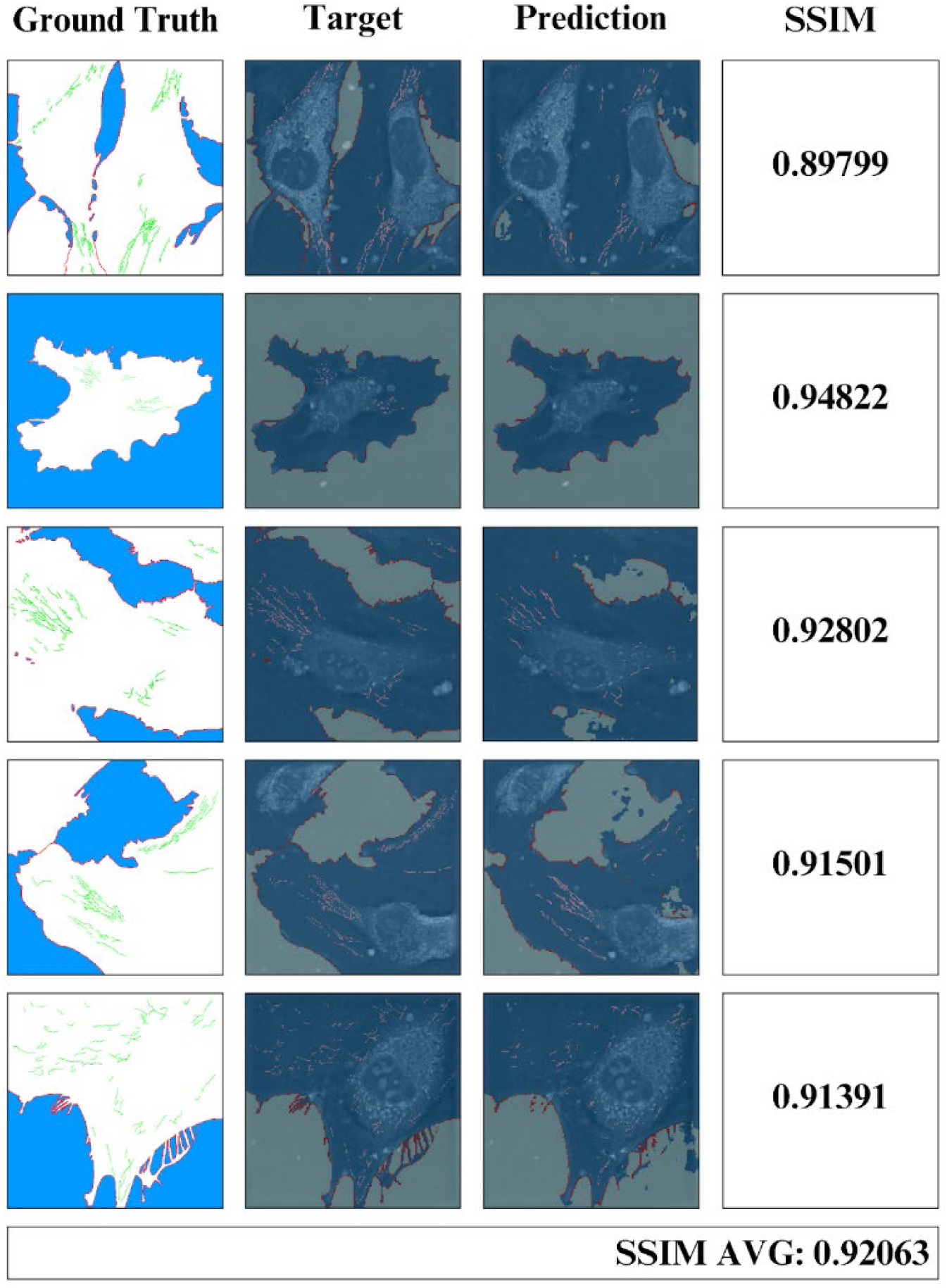
Test dataset consisting of five holotomographic microscopy images and their ground truth, target mask, model prediction, and individual and average Structural Similarity Index for Measuring Image Quality.

To analyze the metrics of the output image from the model, we used current image analysis tools to compare the SSIM values of each output. Figure 6 illustrates the use of ImageJ and CellProfiler tools on a test image to test mitochondrial detection. Comparison of the output of these two tools with the model output image (Figure 6 (a)) indicated substantial variability in the segmentation of each class: mitochondria, cell borders, cytoplasm, and background. To corroborate the visual identification of each desired structure, we calculated the SSIM value for each tool, to compare the ground truth to each output image (values at the bottom of Figure 6). Higher values indicate greater similarity between compared images. The highest values were observed for the model output image, which showed higher similarity to the ground truth than the two existing tools.

**Figure 6.**
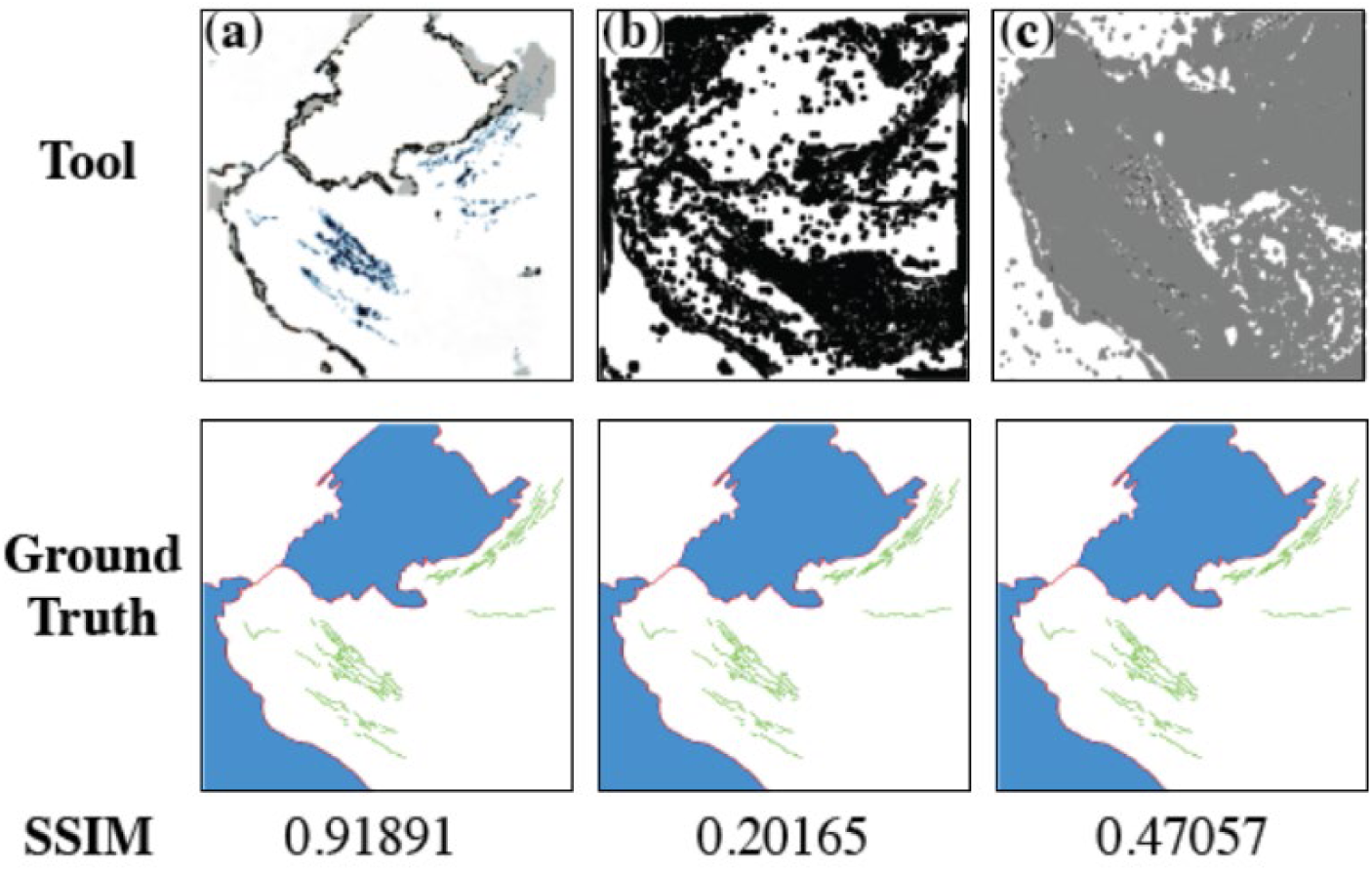
Examples of existing tools compared with ML model output: (**a**) model output image, (**b**) CellProfiler output, and (**c)** ImageJ Mitochondrial Analyzer Plug-in output.

With the CellProfiler coded pathway, the primary and secondary objects were outlined for mitochondria and cell borders, respectively. For the interior and exterior of the cell, a mask was added at the end of the pathway to fill out the remainder of the image. The output image (Figure 6 (b)) shows a general outline of the cell, including darker shadowing of the nucleus and center of the cell. The mitochondrial detection indicates a lack of proper segmentation, observable as dark blending of the cell’s center organelles and the mitochondrial networks. This directly impacted the SSIM value of the image when compared to the actual observation, resulting in a calculated value of 0.20165 or 20.165% similarity between the two images. In addition, CellProfiler’s basic coding block pathway does not enable color assignment for each object.

ImageJ’s Mitochondrial Analyzer Plug-in (Figure 6 (c)) is a pre-coded plug-in that was downloaded to a local computer. The same HTM test image was opened, and the plug-in was selected to automatically analyze the image. Although designated for confocal microscopy, the plug-in was able to fully run the HTM test image. The rightmost image in Figure 6 shows output from ImageJ, indicating successful identification of the cell and general outlining of the mitochondrial networks. Although most were correctly identified, areas such as the top third of the image showed improper pixel assignment and missed areas of the cell border, resulting in an SSIM value of 0.47057 or 47.057% similarity between the two images.

## Conclusions

We have developed a ML algorithm that accurately segments and identifies four classes—cells, cell borders, mitochondria, and background—in HTM images of ECs. The model demonstrated high SSIM, precision, recall, F1-score, and accuracy on an unseen test set, efficiently and effectively segmenting each class within the input images. Improvements over prior models include parameter optimization and the use of a weighted loss function to address overfitting, leading to enhanced model accuracy. The enhanced detection capabilities enable analysis of images with a complete field of view of an entire EC, capturing the full circumference of the cell border and all mitochondrial networks within the cell. This advancement allows for more comprehensive studies of mitochondrial morphology and dynamics in live cells without the need for fluorescent labeling, thereby preserving cell viability and natural behavior.

Future models may incorporate additional classes to identify other subcellular organelles and components with distinct RI signatures, such as lipids or nuclei, further expanding the utility of this approach. By leveraging HTM’s video acquisition capabilities, our method can be extended to track mitochondrial network dynamics over time, providing valuable insights into mitochondrial fusion and fission events in live ECs. This technique holds significant potential for advancing the understanding of EC metabolism and its role in cardiovascular disease progression. By enabling label-free, real-time visualization and quantification of mitochondrial networks, our method may facilitate the development of novel therapeutic strategies targeting mitochondrial dysfunction in ECs.

The availability of our model via a public web application encourages its adoption and validation by the research community, promoting collaborative improvements and broader applications. Future work will focus on increasing the dataset size, incorporating diverse cell types, and enhancing the model’s generalizability to other imaging modalities. In summary, our ML approach represents a significant advancement in the label-free visualization and segmentation of mitochondrial networks in live endothelial cells, with broad implications for cellular biology and disease research.

## Acknowledgements

This study was funded by the National Institute of General Medical Sciences of the National Institutes of Health under Award Number SC2GM140991. The authors thank Clarity in Science Editing and Writing for scientific manuscript editing.

